# Analyses of exon 4a structure reveal unique properties of Big tau

**DOI:** 10.1101/2025.09.04.674244

**Authors:** Itzhak Fischer, Peter W. Baas

## Abstract

Tau is a microtubule-associated protein that modulates the dynamic properties of microtubules and is involved in neurodegenerative diseases known as tauopathies. Tau is expressed as multiple low molecular weight (LMW) isoforms in most neurons of the central nervous system but only as a high molecular weight isoform in neurons of the peripheral nervous system and in a few types of central neurons. Big tau is defined by the inclusion of the alternatively spliced exon 4a, which adds about 250 amino acids to the domain of tau that projects away from the microtubule. Despite low sequence conservation of exon 4a, its length remains remarkably consistent across vertebrates. Here, we analyzed the charge distribution, hydrophobicity, and aggregation propensity of the human sequences of LMW tau, Big tau and the stretch of amino acids encoded by exon 4a. The exon 4a amino acids display a pronounced net negative charge (acidic/basic ratio = 1.30), a consistently hydrophilic composition (average Kyte-Doolittle score = -0.9259) and low β-sheet content of 4.78%. This contrasts with LMW tau, which is more hydrophobic (-0.8930) and contains extended aggregation-prone motifs within the microtubule-binding domain including high β-sheet content of 17.33%. The inclusion of exon 4a in Big tau shifts the global hydrophobicity to intermediate values (-0.9036) and reduces predicted β-sheet content to 13.14%, suggesting decreased aggregation potential. Evolutionary analyses across mammals, birds, and amphibians (human, rat, zebra finch, frog) confirms the minimal sequence identity (16-24% identity in non-mammals) and conserved exon size but show preservation of net negative charge (acidic/basic ratio 1.3-2.3), indicating convergent retention of charge-based properties. Hydrophilicity was also broadly conserved, though less invariant across species. These results demonstrate that exon 4a introduces a highly acidic, hydrophilic module that counterbalances the aggregation-prone domains of LMW tau. The conservation of size and structural properties of the exon-4a-encoded stretch of amino acids, despite sequence divergence, implies strong evolutionary pressure to maintain biophysical properties that counteract pathogenic misfolding.

## INTRODUCTION

Tau proteins are microtubule-associated proteins that regulate cytoskeletal dynamics, axonal transport, and synaptic function (Baas and Qiang, 2019; Alonso et al., 2024). Their pathological aggregation into abnormal filaments is a defining feature of several neurodegenerative diseases collectively termed tauopathies (Avila, 2006; Wang and Mandelkow, 2016). The aggregation process of tau is complex and includes not only aggregation-prone domains of the protein, but is also driven by mutations, posttranslational modifications, cellular stress factors and tau fibrils acting as seeds (Limorenko and Lashuel, 2021).

Among tau isoforms, Big tau is distinguished by the inclusion of exon 4a, an alternative exon encoding ∼250 amino acids (Goedert et al., 1992; Boyne et al., 1995; Fischer and Baas, 2020). The tau 4a exon distinguishes Big tau from its lower molecular weight (LMW) counterparts, but the resulting alterations in tau function have remained enigmatic since its discovery in the early 1990s. What is established is that the large size of the stretch of approximately 250 amino acids encoded by exon 4a (hereafter referred to as exon 4a protein) dramatically extends the projection domain and increases the overall size of the tau protein by nearly 60% compared to LMW tau. This major extension is likely to affect both the structural and functional properties of tau. Indeed, Big tau is selectively expressed in the peripheral and autonomic nervous systems, as well as in distinct regions of the central nervous system such as the cerebellum and brainstem (Boyne et al., 1995; Fischer, 2024).

Surprisingly, the amino acid sequence of the exon 4a protein shows very low conservation, even among mammals (e.g., only ∼50% sequence identity between primates and rodents), and almost no sequence identity with non-mammalian vertebrates (birds, reptiles, amphibians, fish). Nevertheless, the size of the 4a insert remains nearly invariant (e.g., 252 residues in humans and 257 in zebra finch). This contrasts sharply with the remainder of the tau protein, where both the N- and C-termini are highly conserved and the microtubule-binding domain (MTBD; encoded by exons 9-12) maintains strong sequence identity across vertebrate species (e.g., ∼95% between human and zebra finch) (Fischer, 2022).

To investigate the structural properties of Big tau relative to LMW tau (human), and to better understand the unusual size conservation of a protein domain with little sequence homology, we analyzed and compared the two proteins with respect to charge distribution, hydrophobicity and aggregation propensity using a variety of computational tools (Housmans et al., 2023). Our goals were: (1) to determine the effect of exon 4a on tau structure (e.g., folding and aggregation) and (2) to assess these properties in an evolutionary context.

Our analyses revealed that exon 4a protein is characterized by a higher overall negative charge than the rest of tau (which contains a greater proportion of positive charges) and is more hydrophilic than LMW tau (which is overall more hydrophobic). These properties suggest that, while LMW tau is prone to aggregation through misfolding and aggregation mediated hydrophobic interactions, the exon 4a protein is relatively resistant to such aggregation. Importantly, incorporation of exon 4a into LMW tau to form Big tau shifts the hydrophobic properties of LMW tau toward greater hydrophilicity, with values intermediate between LMW tau and exon 4a protein. Analysis of the secondary structure verifies the potential effects of inclusion of 4a exon on aggregation, showing decrease in β strands. This likely reflects the strong influence of the ∼250 amino acid exon 4a insertion on the overall structural profile of Big tau. Consequently, the misfolding and aggregation potential of Big tau appears to be reduced. Another key finding is that the negative charge properties of exon 4a are conserved across species, while the hydrophilic properties vary. Despite low sequence conservation, the size and the structural feature of net negative charge is consistently found in different vertebrate species from amphibians to mammals, which often maintain a hydrophilic structure, suggesting evolutionary pressure to maintain these properties that ultimately define Big tau.

## METHODS

1. **Protein sequences:** Human tau isoforms and exon 4a sequences were retrieved from UniPro and Ensembl as previously described (Fischer, 2022). Comparative sequences were obtained for rat, zebra finch, and frog as follows: **Homo sapiens** microtubule-associated protein tau (MAPT) ENST00000415613.6 MAPT-205 UniPort:P10636-9 EPESGKVVQEGFLREPGPPGLSHQLMSGMPGAPLLPEGPREATRQPSGTGPEDTEGGRHAPELLKHQLLGDLHQEGPPLKGAGGKERPGSKEEVDEDRDVDESSPQDSPPSKASPAQDGRPPQTAAREATSIPGFPAEGAIPLPVDFLSKVSTEIPASEPDGPSVGRAKGQDAPLEFTFHVEITPNVQKEQAHSEEHLGRAAFPGAPGEGPEARGPSLGEDTKEADLPEPSEKQPAAAPRGKPVSRVPQLK **Rat** Rattus norvegicus ENSRNOT00000042984.6, MAPT-207 UniPort: F1LST4 EPQKVEIFSQSLLVEPGRREGQAPDSGISDWTHQQVPSMSGAPLPPQGLREATHQPLGTRPEDVERSHPASELLWQESPQKEAWGKDRLGSEEEVDEDITMDESSQESPPSQASLAPGTATPQARSVSASGVSGETTSIPGFPAEGSIPLPADFFSKVSAETQASPPEGPGTGPSEEGHEAAPEFTFHVEIKASAPKEQDLEGATVVGAPAEEQKARGPSVGKGTKEASLLEPTDKQPAAGLPGRPVSRVPQLK **Zebra Finch** ENSTGUT00000021320.1 MAPT-208 GEPSSPKLQPGPRERVGEAVKSASQPPEQGLGPQQPPLSRETKAPAAAPTRIEVTIPIPLDMYQGSEGSGELWDQGGTEGLARAGGTGGHKDGPSPLCARATIKEDSGGRERDEDRDIDETSGQGLPSLVDQCVSLAPEGSCPAAAQEAREEYDGENKSKGVLRDTPGEALLVEAESHKAGEDQEEKRELLEGEGGPDSALSEPSGSVSLKEAEPREGEDSGPVLETAKLPAEGEDGVKKVDEDAPVGEAVPDAGGRRTPRRKPGGLAADKASRVPLLK **Frog** tropical clawed frog, Xenopus tropical ENSXETT00000084149.2 MAPT-206 UniPort: A0A6I8RGV8 EEIALLAAAGQEEEYEMDTMEETLKITAKDQTHAENYGITGDVDGESQNDETALSSGMVESAV EEDYYKETNGKEVNLEICEDDTEGWEEQIDEGIIMQDSVAPPKGGEQELSSVEQPQTNGTGAEHI FLEDNQHKKDTEEPFMAIPANSFPVGQIRPRASVSVYQVEI DANIPIDSKEAPCEDVGIPGGTKVDTERATEETLKSPRKRMPAHGSGIPVSRVPVPK
2. **Charge distribution:** Acidic and basic residues were quantified, and cumulative charge profiles were plotted. Ratios of acidic to basic residues were calculated per domain.
3. **Hydrophobicity analysis:** The Kyte-Doolittle plot is a simple but powerful method to visualize hydrophobic vs hydrophilic regions of a protein sequence, providing insights into folding, solubility, membrane-association, and aggregation (Kyte and Doolittle, 1982). After applying it to LMW tau, exon 4a protein, and Big tau, average hydrophobicity values were calculated.
4. **Aggregation propensity:** Secondary structure and β-sheet formation were predicted with PASTA 2.0. Amyloid-prone regions were identified with AggreProt and Aggrescan. **PASTA 2** is an energetic function derived from high-resolution protein structures, which considers interaction potential and H-bond formation between all non-consecutive residues for parallel and anti-parallel β -pairing (Walsh et al., 2014).
5. **AggreProt**, a machine learning sequence-based predictor of Amyloid-Prone Regions (APRs), was designed to detect short, amyloid-related and biologically relevant APRs, no longer than 50 residues (Planas-Iglesias et al., 2024). **Aggrescan** prediction is assayed against an aggregation propensity scale for the 20 proteinogenic amino acids derived from in vivo experiments (Conchillo-Sole et al., 2007).
6. **Tango w**as used to confirm the biophysical properties of the exon 4a protein, specifically its charge and hydrophilicity - http://tango.crg.es/
7. **Evolutionary analysis:** Pairwise sequence identity was determined with Clustal Omega. Charge and hydrophobicity profiles were compared across species.

## RESULTS

### 1. Comparing LMW Tau and Exon 4a (Human)

#### 1a. Charge distribution

We examined the charge distribution, which is a critical determinant of protein structure, function, and molecular interactions. As shown in Fig. 1, LMW tau contains a relatively balanced number of acidic and basic residues, with a slight excess of negative residues. As shown in Fig. 1, charge distribution varies across domains. The exon-4a protein shows a significantly higher number of acidic residues compared to basic ones, resulting in a strong overall negative charge. The ratio of negative to positive residues underscores this difference: LMW tau: 61 negative / 55 positive (ratio = 1.11). Exon 4a: 43 negative / 33 positive (ratio = 1.30).

**Fig. 1.**
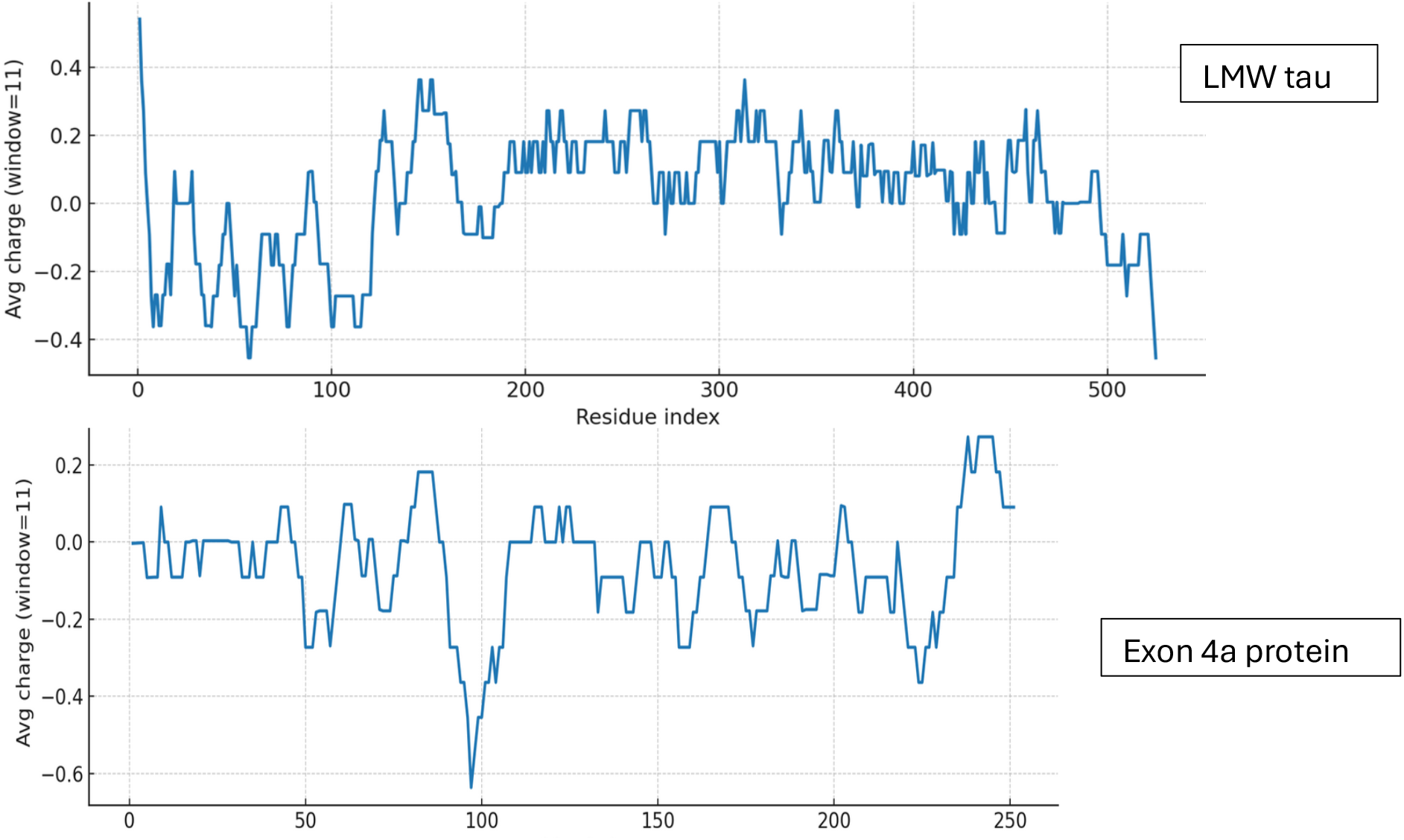
Charge distribution of LMW tau and exon 4a protein.

### 1b. Hydrophobicity Profile

The distribution of hydrophobic versus hydrophilic residues strongly influences protein folding and structural stability. Using the Kyte-Doolittle scale, Fig. 2 illustrates average hydrophobicity values. Scores closer to zero or positive reflect hydrophobicity, while more negative scores indicate hydrophilicity. As shown in Fig. 2, LMW tau displays a mixed profile, with both hydrophobic and hydrophilic regions. Several peaks above the zero line reflect hydrophobic stretches that could form buried cores or, if sufficiently long, potential transmembrane segments. Its elevated glycine and proline content suggests greater structural flexibility and intrinsically disordered regions (IDR). The exon 4a protein remains consistently below zero, with only minimal hydrophobic peaks. This indicates a predominantly hydrophilic nature, consistent with its strong negative charge, suggesting that exon 4a is a soluble protein localized to aqueous environments or exposed surfaces of larger complexes. The average values for are for LMW tau -0.8930 and for exon 4a protein: -0.925. Note the “hot spots” of aggregation in the LMW tau shown in more details in Table 1.

**Table 1.** Aggregation analysis with sequence features for the different domains of LMW tau

| <u>Region</u> | <u>Sequence features</u> | <u>Aggregation tendency</u> |
| --- | --- | --- |
| **N-terminal (1-150)** | Acidic, Pro/Gly-rich | **Low**, mostly disordered, soluble |
| **Projection domain (151-244)** | Many PXXP motifs, positive charges | **Low-moderate**, can regulate conformational transitions |
| **NTBD (245-368)** | Contains PHF6* and PHF6 | **Very high**, nucleation core |
| **C-terminal (369-441)** | Acidic, charged, extended | **Moderate-low**, stabilizes but not strongly amyloidogenic |

**Fig. 2.**
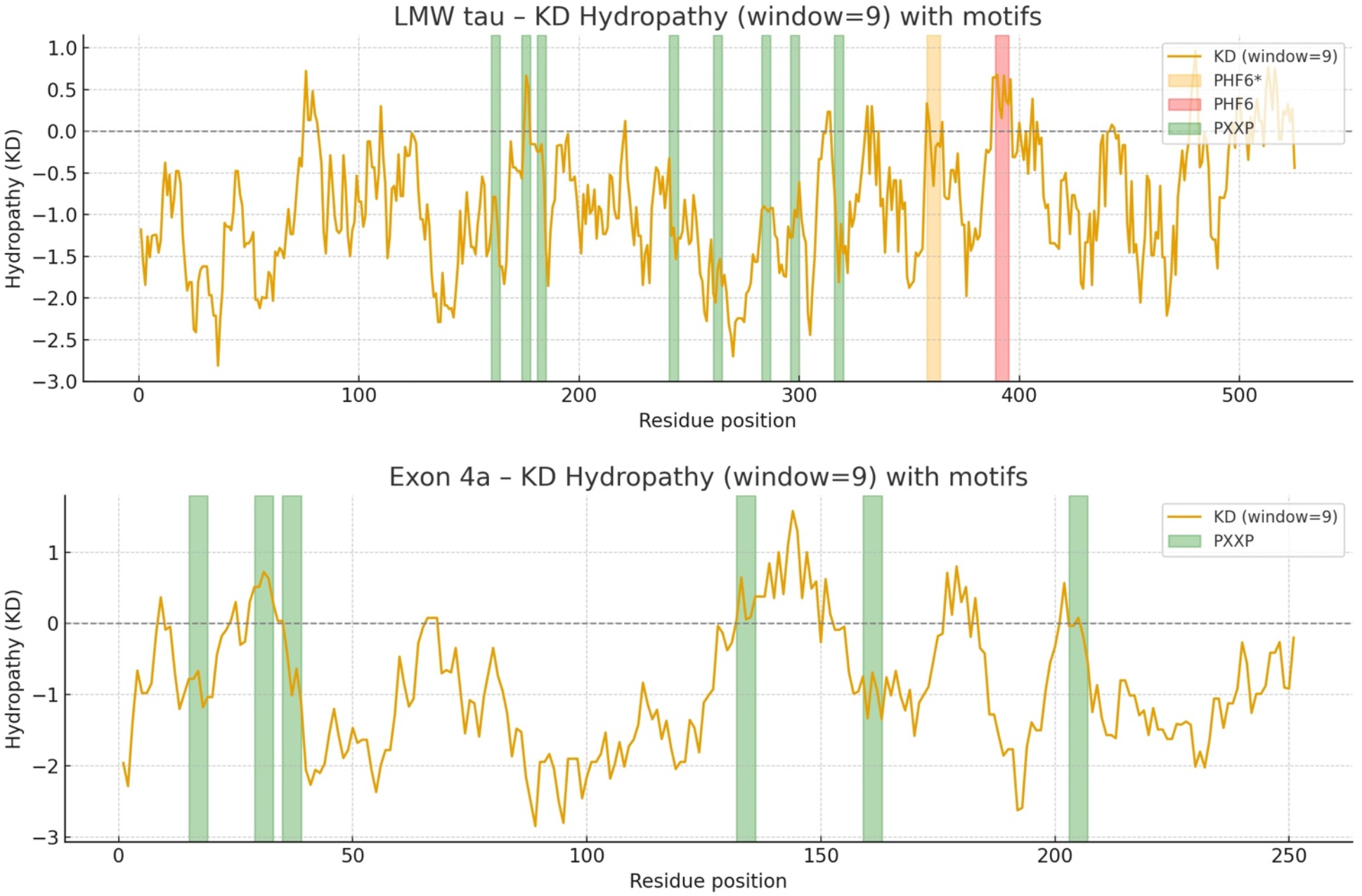
The Kyte-Doolittle hydrophobicity plot annotated with PHF6* and PHF6 as key aggregation “hot spots” and PXXP motifs as potential binding partners and regulatory interactions.

### 1c. Analysis of different LMW tau domains

To obtain more detailed information than the average values of LMW tau we analyzed the 3 major domains of the protein as shown in Fig. 3 for the ratio of negative to positives residues in Panel A and hydrophobicity values in Panel B.

**Fig. 3.**
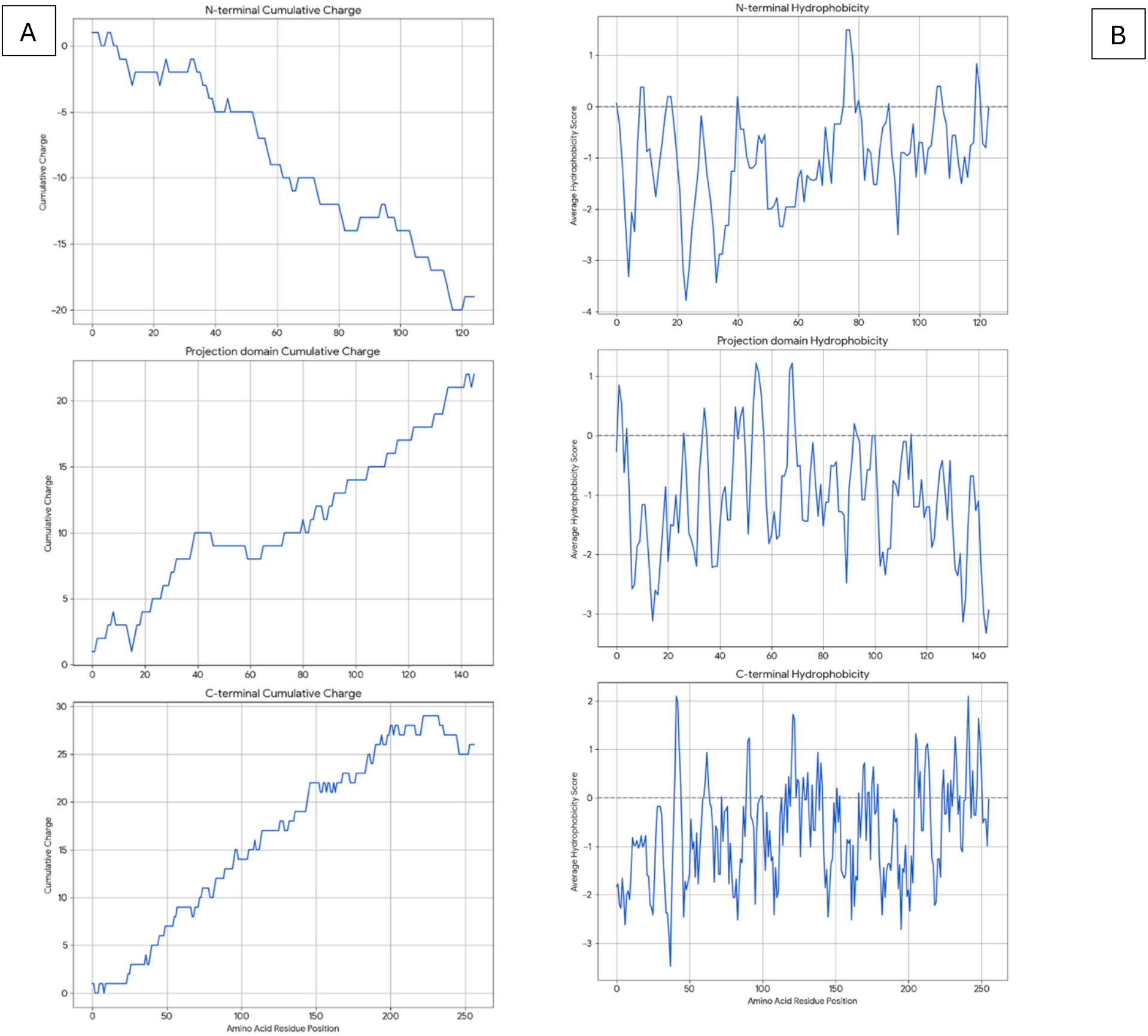
Analysis of LMW tau domains for charge distribution (Panel A) and hydrophobicity (Panel B) 1d. Analysis of LMW tau Aggregation

**Fig. 4.**
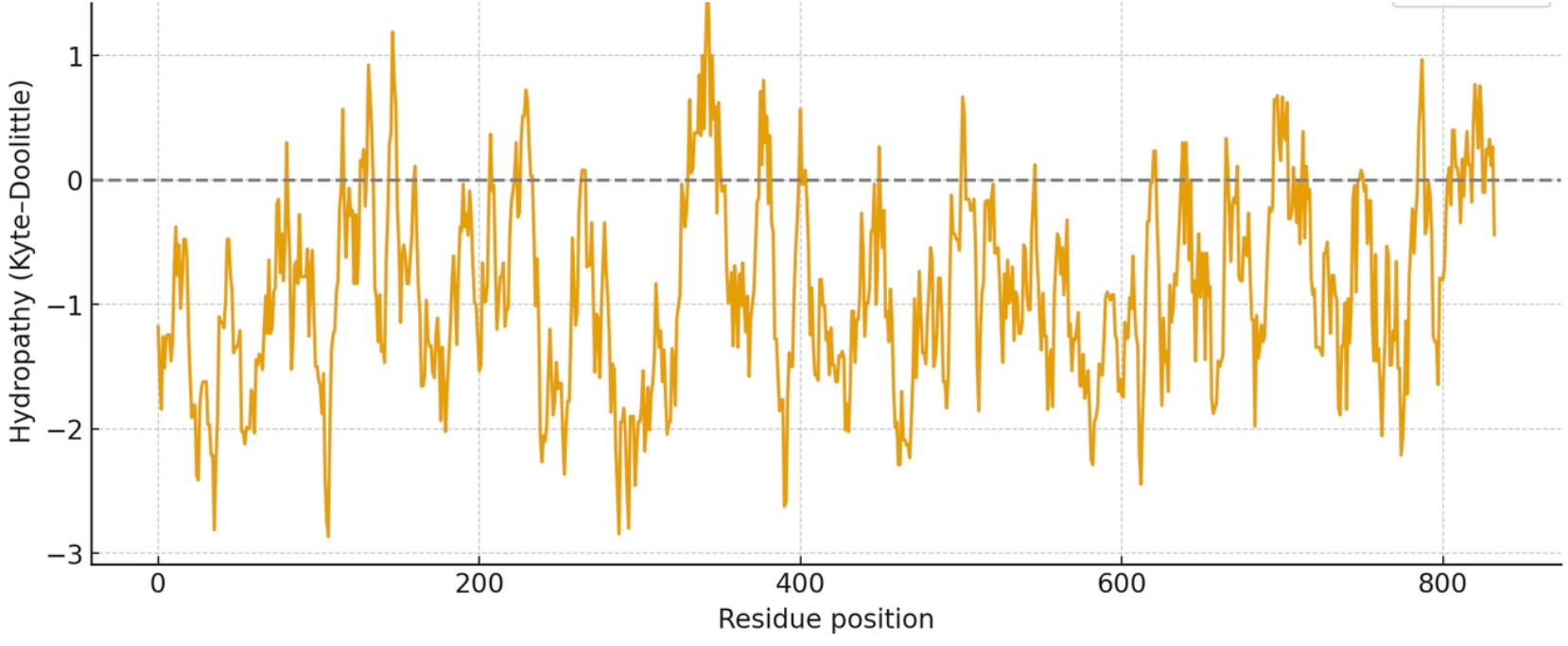
Hydrophobicity plot of Big tau (LMW tau + exon 4a)

#### N-terminal

The cumulative charge plot for the N-terminal protein shows a steep and consistent downward trend. This confirms its highly acidic nature, as negative charges accumulate rapidly along the sequence with an average ratio of 3. The hydrophobicity plot for this protein remains consistently in the negative range, indicating a highly hydrophilic and water-soluble peptide with a value of -1.0573. The fluctuations are minimal, which is typical for proteins that are largely disordered.

#### Projection Domain

The cumulative charge plot for the projection domain shows a steep and consistent upward trend. This reflects its highly basic nature, as positive charges accumulate along the sequence resulting in a low ratio of 0.28. Like the N-terminal peptide, this domain is also highly hydrophilic, with its hydrophobicity plot remaining consistently in the negative range at -1.0717. Its average hydrophobicity is the lowest of the three, making it the most hydrophilic overall.

#### C-terminal

The cumulative charge plot for the C-terminal protein shows a steep and consistent upward trend. This reflects its highly basic nature, as positive charges accumulate along the sequence resulting in a low value of 0.47. While still in the hydrophilic range, this protein is significantly less hydrophilic than the N-terminal protein with a value of -0.7121. The plot shows several distinct hydrophobic peaks that rise closer to the zero line, suggesting the presence of more hydrophobic regions. These regions could be involved in protein folding or interactions with other molecules.

The aggregation propensity of LMW tau was examined using the AggreProt program which selected four different domains shown in Table 1. It highlights the known high aggregation domain of the MTBD repeats, which include the PHF6 motifs.(Ganguly et al., 2015).

2. Comparing Hydrophobicity of LMW Tau and Big Tau. PXXP motifs can be protein–protein interaction domains with SH3 containing partners, role in signaling and cytoskeletal regulation. The various PHF6 domains drive abnormal aggregation into paired helical filaments (PHFs) hallmark of Alzheimer’s disease and other tauopathies.

### 2a. Hydrophobicity

Given the marked structural differences between LMW tau and exon 4a, we next examined Big tau, which incorporates the exon 4a sequence. Hydrophobicity analysis revealed that Big tau shows intermediate hydrophilic values between LMW tau and exon 4a, reflecting the substantial influence of the exon 4a domain. Thus, with the value of LMW tau at -0.8930 and exon 4a at -0.9259 the combined hydrophobicity value of Big tau showed at -0.9036 which is the average value between LMW tau and exon 4a: -0.9094. On the Kyte–Doolittle scale (Fig. 3), this shift is most evident between residues 120-375, corresponding to the exon 4a region. These values confirm that inclusion of exon 4a alters the global hydrophobicity of tau, reducing the overall hydrophobic properties of tau.

2 b. Analysis of aggregation propensity.

Using PASTA 2 analysis we determined the secondary structure of the protein with focus on β-sheet– forming sequences are the primary drivers of aggregation. Once formed, β-sheets template further misfolding, creating a self-propagating process. Mutations that increase β-sheet propensity (e.g., replacing polar residues with hydrophobic ones) usually enhance aggregation. Exon 4a showed a much the lower values for β-sheet relative to LWM tau (4.78 vs 17.33), which resulted in Big tau a lower value, a decrease of almost 25% (13.14 vs 17.33). Interestingly, the exon 4a-L had a higher value of β-sheet relative to 4a suggesting that it may not be as effective in reducing the aggregation potential.

### 2c. Aggregation analysis using AggrePlot

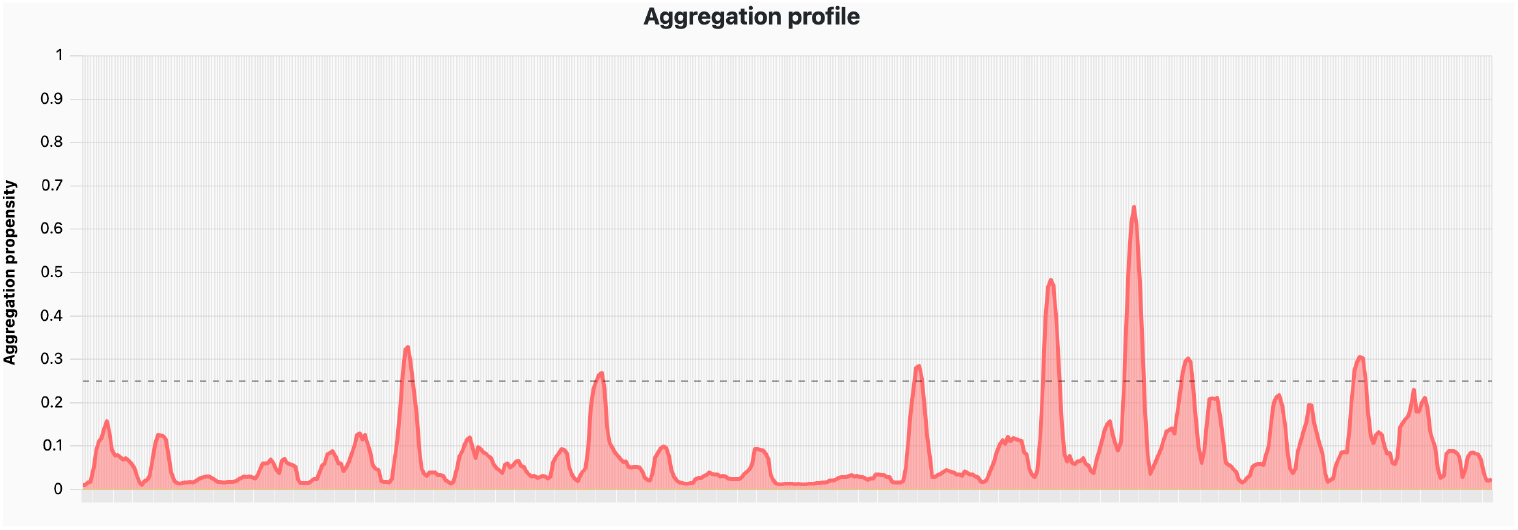

**LMW tau**

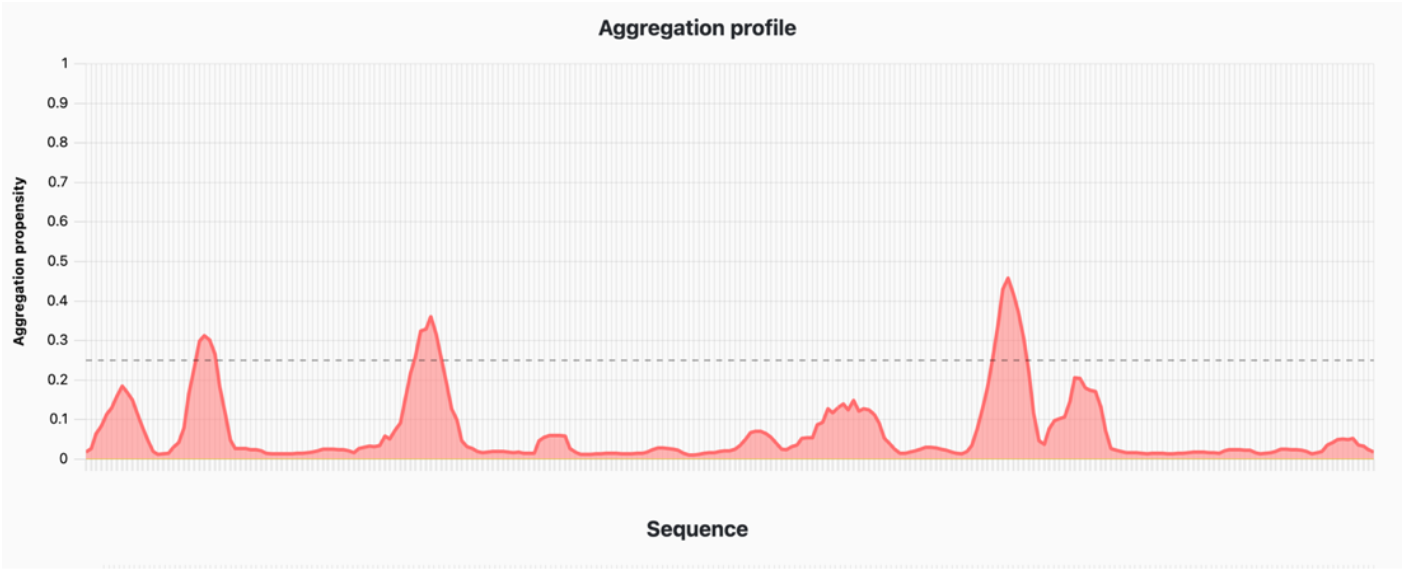

**Exon 4a**

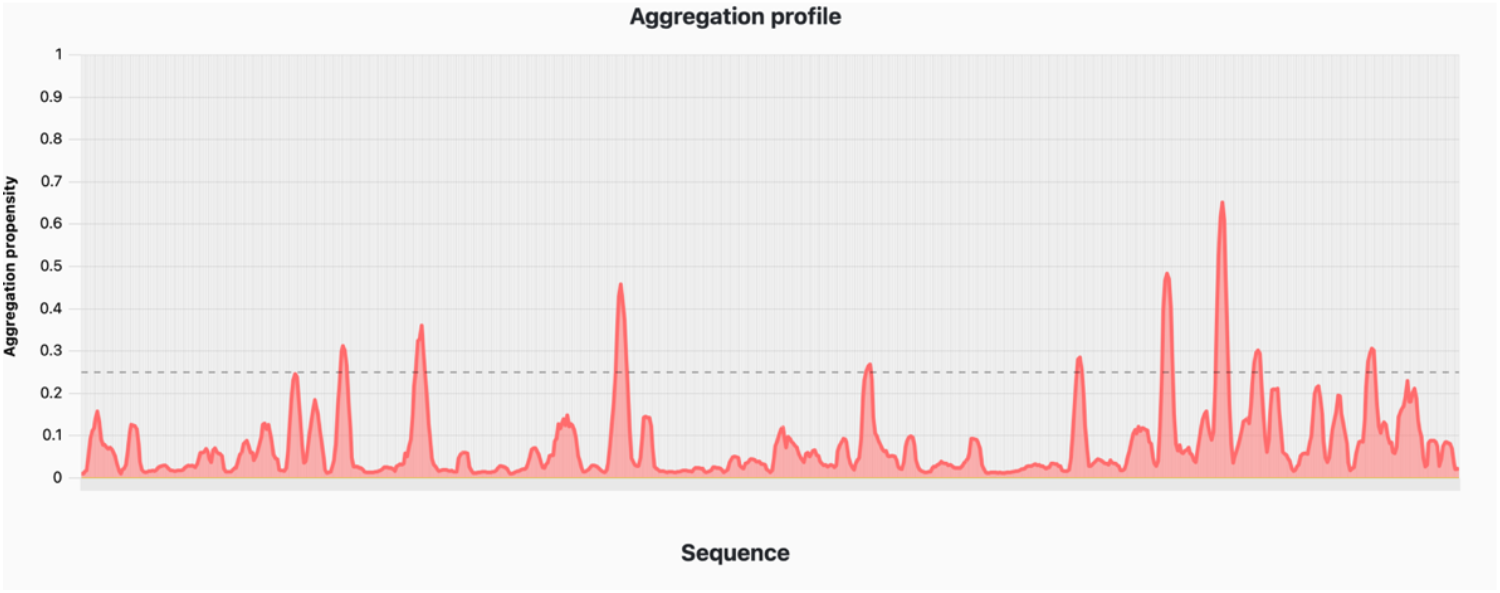

**Big tau**

## 3. Evolutionary Perspective

Previous studies have shown that the amino acid sequence of the exon 4a protein shows very low conservation, even among mammals (e.g., only ∼50% sequence identity between primates and rodents), and almost no sequence identity with non-mammalian vertebrates (birds, reptiles, amphibians, fish). Nevertheless, the size of the 4a protein remains similar (ranging in the examples below 226-280). This contrasts sharply with the remainder of the tau protein, where both the N- and C-termini containing the MTBD maintains strong sequence identity across vertebrate species as shown in Table 2.

**Table 2.** Secondary structure analysis PASTA 2 with emphasis on % β-strand.

| Protein | Length | # Amyloids | Best Energy | % Disorder | % α-helix | % β-strand | % Coil |
| --- | --- | --- | --- | --- | --- | --- | --- |
| Exon 4a | 251 | 0 | -4.10 | 96.01 | 11.95 | 4.78 | 83.27 |
| LMW tau | 525 | 1 | -5.51 | 68.19 | 6.86 | 17.33 | 75.81 |
| Big tau | 776 | 1 | -5.51 | 78.47 | 8.25 | 13.14 | 78.61 |
| Exon 4a-L | 355 | 0. | -4.10. | 91.54 | 7.89. | 6.2 | 85.92 |

**Table 2.** Analysis of sequence identity across vertebrates

|  | Human | Rat | Zebra finch | Frog |
| --- | --- | --- | --- | --- |
| LMW tau | 100 | 71 | 52 | 43 |
| N-terminal | 100 | 83 | 60 | 38 |
| C-terminal | 100 | 97 | 91 | 80 |
| 4 exon protein | 100 | 56 | 24 | 16 |

To assess whether the unique properties of human exon 4a protein are conserved across vertebrates, we analyzed charge and hydrophobicity profiles of exon 4a protein in representative species:

Negative to positive charge ratio relative to LMW tau at 1.1

### Exon 4a

- Human: 1.3
- Rat: 1.58
- Zebra Finch: 1.56
- Frog: 2.32

Average hydrophobicity scores relative to LMW tau at -0.8930

### Exon 4a

- Human: -0.9259
- Rat: -0.7677
- Cat: -0.9556
- Zebra Finch: -0.9061
- Frog: -0.7770

All of these exon 4a proteins displayed negative scores, confirming hydrophilicity as a conserved feature. Proteins with less negative values (rat and frog) exhibited more peaks approaching the zero line, reflecting localized hydrophobic stretches.Taken together, these results indicate that, despite low sequence conservation, exon 4a protein maintains consistent structural properties across vertebrate evolution: stable size, net negative charge, and hydrophilicity. These conserved features likely serve to mitigate tau misfolding and aggregation.

## DISCUSSION

This study describes structural and evolutionary analyses of tau’s exon 4a, the defining feature of the high molecular weight Big tau protein (Fischer and Baas, 2020). Our findings address the longstanding question of how a protein domain with minimal sequence conservation can nonetheless maintain a consistent length and functional role across diverse vertebrate lineages (Fischer, 2022). Using computational tools which are sequence-based and structure-based such charge distribution, Kyte-Doolittle plot, PASTA 2.0, AggreProt Aggrescan and Tango (Navarro and Ventura, 2022; Housmans et al., 2023), we have shown that although the primary amino acid sequence encoded by exon 4a is highly divergent, its key biophysical properties including charge distribution, hydrophobicity, and aggregation profile are conserved. These features counterbalance the inherent aggregation propensity of LMW tau, suggesting that exon 4a represents an evolutionary innovation to mitigate tau misfolding. Specifically, our analyses indicate that inclusion of exon 4a modifies the structural properties of tau by introducing a highly acidic, hydrophilic, and intrinsically disordered domain. Given that protein misfolding and aggregation are frequently driven by exposed hydrophobic patches, the insertion of such a hydrophilic, negatively-charged domain likely reduces intermolecular association by increasing solvation and electrostatic repulsion (Levine et al., 2015; Hernandez et al., 2022). This also effectively reduces the β-sheet secondary structure and aggregation potential. Interestingly, exon 4a-L may not be as effective as 4a: Its average hydrophobicity is -0.8794 vs -0.9259), which is similar to LMW tau, and it has a higher value of β-sheet than 4a suggesting that it may not be as effective in reducing the aggregation potential and thus play a different role possibly related to its larger size of 330 amino acids. Indeed, the 4a-L was found in prostate cancer cell lines where microtubules are mostly associated with cell division (Souter and Lee, 2010). We propose a model in which the large exon 4a-derived stretch of about 250 amino acids increases dramatically the projection domain and together with its unique biophysical properties distinguishes the Big tau protein from the LMW tau. These modifications likely affect the physiological roles of Big as well as attenuate pathological misfolding, in region of the nervous system with high expression of Big tau.

### Functional implications

The proposed model aligns with observations that Big tau is selectively expressed in neurons of the peripheral and autonomic nervous systems, as well as in specific CNS regions such as brainstem and cerebellum, which are typically less vulnerable to tauopathies (Boyne et al., 1995;Fischer and Baas, 2020;Chung et al., 2024). The biophysical properties of exon 4a protein may therefore represent a molecular adaptation to enhance proteostatic resilience in these neuronal populations, which must maintain long axons and high transport demands over the lifespan. This model also aligns with emerging therapeutic strategies aimed at modulating tau aggregation and interfering with seed propagation.

Beyond its role in aggregation suppression, Big tau is also likely to affect physiological properties such as microtubule dynamics, axonal transport, and interactions with cytoskeletal proteins (Fischer, 2022). The expanded projection domain could increase inter-microtubule spacing, reduce crosslinking density, and alter bundling properties, thereby creating a cytoskeletal architecture optimized for long-range axonal viability. Moreover, by decreasing the density of tau molecules bound to microtubules, Big tau may reduce steric hindrance for motor proteins, generating a more permissive substrate for kinesin- and dynein-mediated transport. Whether these effects are mediated solely by the size of the exon 4a extension, or also by its specific physicochemical properties, will require exploration in experimental cell systems.

### Evolutionary perspective

Comparative analyses of human, rat, zebra finch, and frog tau reveal a striking pattern where exon 4a shows extensive primary-sequence divergence yet contributes a highly conserved length and biophysical profile to the tau protein. This combination of low sequence identity with conserved composition is characteristic of IDR, which are often under selection for ensemble properties (e.g., charge density, hydrophobicity, proline/glycine content) rather than sequence motifs (Zarin et al., 2017; Holehouse and Kragelund, 2024; Singleton and Eisen, 2024). Our hydrophobicity analyses reinforce this view, showing that exon 4a protein is more hydrophilic than LMW tau, while Big tau as a whole occupies an intermediate state consistent with evolutionary pressure to preserve solubility and suppress β-sheet formation within the expanded projection domain. By contrast, the MTBD is highly conserved across species and contains short hexapeptides that drive aggregation (Ganguly et al., 2015). Exon 4a protein appears to act as a solubilizing, charge-rich spacer that offsets the aggregation risk conferred by the MTBD. This may be especially advantageous in peripheral nervous system neurons, which face extreme transport distances and high molecular crowding along axons.

The near-fixed length of exon 4a across birds, amphibians, and mammals suggests geometric or structural constraints: a shorter insertion might fail to provide sufficient solubilizing capacity, while a longer one could impose steric or metabolic costs. The convergent retention of charge density and hydrophilicity, despite sequence divergence, indicates that selection has operated on physicochemical properties rather than precise sequence motifs. This modular organization of conserved functional “core” domains (e.g., MTBD, C-terminal tail) combined with compositionally conserved IDR “spacers” underscores how tau balances structural versatility with proteostatic resilience. Although all the tested species maintain an overall hydrophilic, acidic profile in exon 4a, our analyses suggest that frogs and rats exhibit somewhat lower net negative charge and more local hydrophobic patches compared to humans and birds. This predicts subtle species-specific differences in chain compaction under physiological ionic conditions. Nonetheless, the preserved length and high overall charge imply that the fundamental role of exon 4a of aggregation suppression and structural spacing is conserved.

From an evolutionary standpoint, exon 4a represents a proteostatic adaptation that buffers the aggregation-prone MTBD, particularly in neurons with long axons and high transport loads. This stabilizing, disorder-based module provides a blueprint for therapeutic design: acidic IDR-like extensions or charge-enhancing modifications grafted onto LMW tau could emulate Big tau’s resistance to misfolding without disrupting MT binding.

### Limitations and future directions

Our sequence-based and bioinformatic analyses provide a strong mechanistic framework, but experimental validation will be essential. *In vitro* aggregation assays, cellular expression systems, and *in vivo* models should be employed to test whether Big tau indeed exhibits lower aggregation kinetics and reduced pathology spread. Furthermore, the roles of post-translational modifications and cell type– specific expression patterns need to be explored to assess whether exon 4a inclusion correlates with functional resilience in vulnerable neuronal populations. Such studies will clarify whether Big tau functions primarily as a structural spacer, a proteostatic buffer, or both.

## Conclusion

Exon 4a defines the unique properties of Big tau by introducing a large, acidic, and hydrophilic domain that reduces aggregation propensity. Despite extensive sequence divergence, the conservation of exon 4a length and charge distribution across vertebrates underscores its evolutionary significance. By conferring resistance to misfolding, Big tau may help explain the relative resilience of peripheral and cerebellar neurons to tau-related neurodegeneration.

## ACKNOWLEDGEMENTS

We acknowledge the online use of computational resources for aggregation predictions detailed in Methods. Thqs work was supported by grants from the USA Natqonal Instqtutes of Health (R21AG068597 and R01NS28785) and the USA Department of Defense (W81XWH2110189) to PWB.

